# Anti-amyloid antibody equilibrium binding to Aβ aggregates from human Alzheimer disease brain

**DOI:** 10.1101/2025.05.20.654902

**Authors:** P. Monroe Butler, Anna Francis, Angela L. Meunier, Amirah K. Anderson, Elizabeth L. Hennessey, Michael B. Miller, Cynthia A. Lemere, Dennis J. Selkoe, Andrew M. Stern

## Abstract

**Importance:** Anti-amyloid immunotherapy is used to treat Alzheimer disease (AD) with moderate benefits and potentially serious side effects due to amyloid related imaging abnormality with effusions/edema (ARIA-E). Different anti-amyloid antibodies have different *in vitro* binding characteristics to different synthetic Aβ aggregates, leading to the assumption that they bind different species in the human brain. Lecanemab is hypothesized to bind “protofibrils,” but these are not well-characterized in human brain. It is also unknown how binding differences correlate with ARIA-E rates. The *APOE ε4* allele increases ARIA-E risk, but how it affects antibody binding characteristics is unknown.

**Objectives:** To determine whether anti-amyloid antibodies bind different species of human brain Aβ and whether these binding properties to human brain Aβ explains ARIA-E rates.

**Design:** Cross-sectional study of 18 postmortem human brains.

**Setting:** Single tertiary care hospital.

**Participants:** Deceased patients with AD and cerebral amyloid angiopathy (CAA).

**Main Outcomes and Measures:** Equilibrium binding constants (K_D_) and total Aβ binding (B_max_) of recombinant aducanumab, lecanemab, and donanemab equivalents to human brain soluble and insoluble amyloid plaque-enriched and CAA-enriched Aβ aggregates.

**Results:** Lecanemab did not bind with greater affinity to the soluble fraction of Aβ compared to aducanumab. All three antibodies were bound essentially identical quantities of Aβ across the 18 cases and fractions (Pearson’s r 0.84 – 0.97). Antibody preference for plaque *vs* CAA Aβ did not differ in soluble fractions but differed slightly in insoluble extracts. The *APOE ε4* allele led to a more soluble antibody-accessible Aβ pool in a dose-dependent manner for all three antibodies.

**Conclusions and Relevance:** The lecanemab binding target in human brain is unlikely to be distinctly “protofibrillar” compared to other antibodies. Differences in antibody preference for plaque *vs* CAA Aβ are unlikely to fully explain differences in ARIA-E rates. The *APOE ε4* allele may plausibly increase ARIA-E risk by making antibody-accessible Aβ more soluble. These results have implications for improving the safety and efficacy of current and future anti-amyloid antibody therapies.

## Introduction

The introduction of disease-modifying therapy is changing the standard of care for Alzheimer disease. Three anti-amyloid antibodies, aducanumab, lecanemab, and donanemab, have received United States Food and Drug Administration approval to treat early Alzheimer disease (AD). Lecanemab and donanemab have entered clinical use. All three antibodies target misfolded, aggregated Aβ, the principal component of amyloid plaques. All three clear amyloid plaques as measured by amyloid positron emission tomography (PET) and avoid binding to monomeric Aβ. However, all three are thought to derive their preference for misfolded, aggregated Aβ over monomeric Aβ through different mechanisms. Donanemab binds to pyroglutamate-3 Aβ, a posttranslational modification found in plaques but not newly synthesized Aβ monomers. Lecanemab and aducanumab both bind conformational epitopes of misfolded Aβ. Between lecanemab and aducanumab, lecanemab is purported to have greater specificity for toxic protofibrils, rather than “mature” amyloid fibrils, and this distinction is postulated to account for its better observed efficacy^1^. However, binding preferences of all three antibodies have been published almost entirely using synthetic *in vitro* Aβ assemblies, rather than Aβ aggregates derived from human brain tissue. Structures of *in vitro* amyloids may differ from *in vivo* ones, and binding properties to *in vitro* assemblies do not infer binding properties to those present in the human brain.

All plaque-clearing anti-amyloid antibodies cause amyloid related imaging abnormality with edema/effusions (ARIA-E), a side effect that can rarely be fatal, and the cause of ARIA-E is not currently known. One hypothesis for the cause of ARIA-E is inflammation and blood-brain barrier breakdown due to binding of anti-amyloid antibodies to cerebral amyloid angiopathy (CAA), Aβ aggregates in the tunica media of brain arterioles and small arteries instead of (or in addition to) in the neuropil^2^. Support for this hypothesis comes from the observations that MRI-detectable lobar microhemorrhages, usually caused by CAA in the AD population, increase risk for ARIA-E; that ARIA-E exhibits clinical and radiographic similarity to CAA-related inflammation (CAAri), a sporadic disease possibly mediated by endogenous anti-Aβ autoantibodies; and autopsies of rare fatal ARIA-E cases have demonstrated vasculitis^3,4^.

These observations can be compiled into a theory in which 1) lecanemab, aducanumab, and donanemab bind structurally different classes of Aβ aggregates in human AD brain, and 2) differences in ARIA-E rates (aducanumab > donanemab > lecanemab) are attributable to differences in binding preference to CAA Aβ vs plaque Aβ. In the current study, we sought to test predictions of this theory: 1) certain populations of Aβ aggregates from human AD brain (*i.e.* “protofibrils”) will be accessible to binding to lecanemab but not aducanumab or donanemab, 2) lecanemab will exhibit greater binding affinity to aqueously extracted (“soluble,” “protofibrillar”) Aβ aggregates than aducanumab, and 3) binding preference of all three antibodies for CAA vs plaque Aβ will follow the inverse rank-order of observed ARIA-E rates in phase 3 clinical trials.

## Methods

### Case selection

We selected eighteen cases from the BWH Brain Bank with pathological diagnoses of both AD and CAA as described in Table 1. As expected for a population enriched for both AD and CAA, the *APOE* genotypes were enriched for the *ε4* allele. We extracted Aβ in two ways as described in detail in the Supplementary Information: 1) an aqueous extraction composed of material in the supernatant after homogenization in TBS followed by centrifugation, and 2) an insoluble extraction from the pellet of the first extraction. Separate extracts were prepared from the grey matter of the occipital lobe and from the overlying leptomeninges.

**Table 1.**
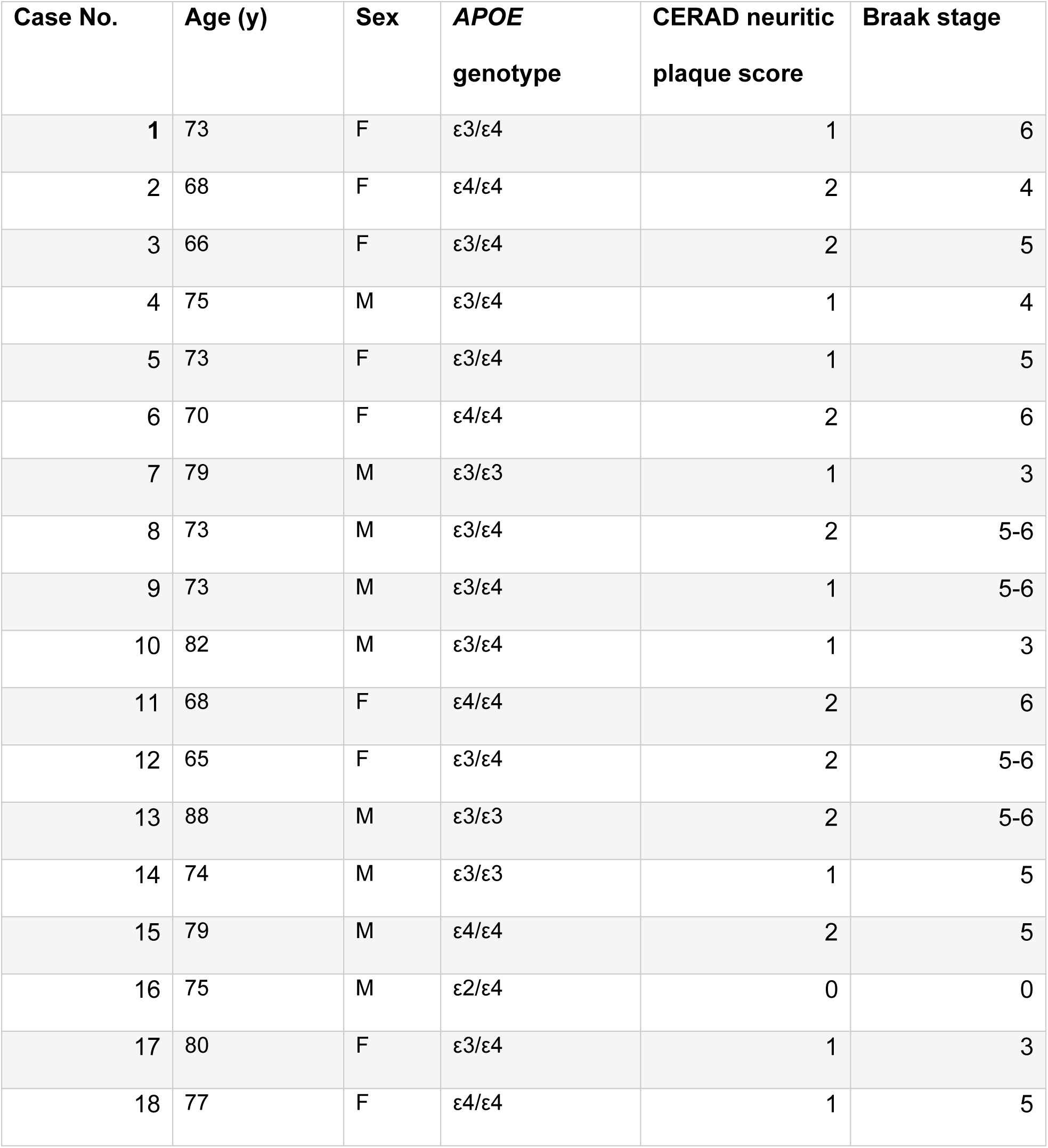
Case characteristics.

### Measurement of equilibrium binding affinity

A full description of the biochemical methods, rationale, and validation is available in the Supplementary Information. Briefly, liquid extracts (aqueously soluble or insoluble, from parenchyma or from meninges) were diluted and immunoprecipitated with serially diluted anti-amyloid antibody on magnetic beads, followed by washing and ELISA quantitation of Aβ42 and Aβ40. The resulting titration curves were fitted to a one-site binding model resulting in a K_D_ (approximation of binding affinity) and B_max_ (total Aβ available to the antibody to bind) in GraphPad Prism software (Fig 1).

**Figure 1.**
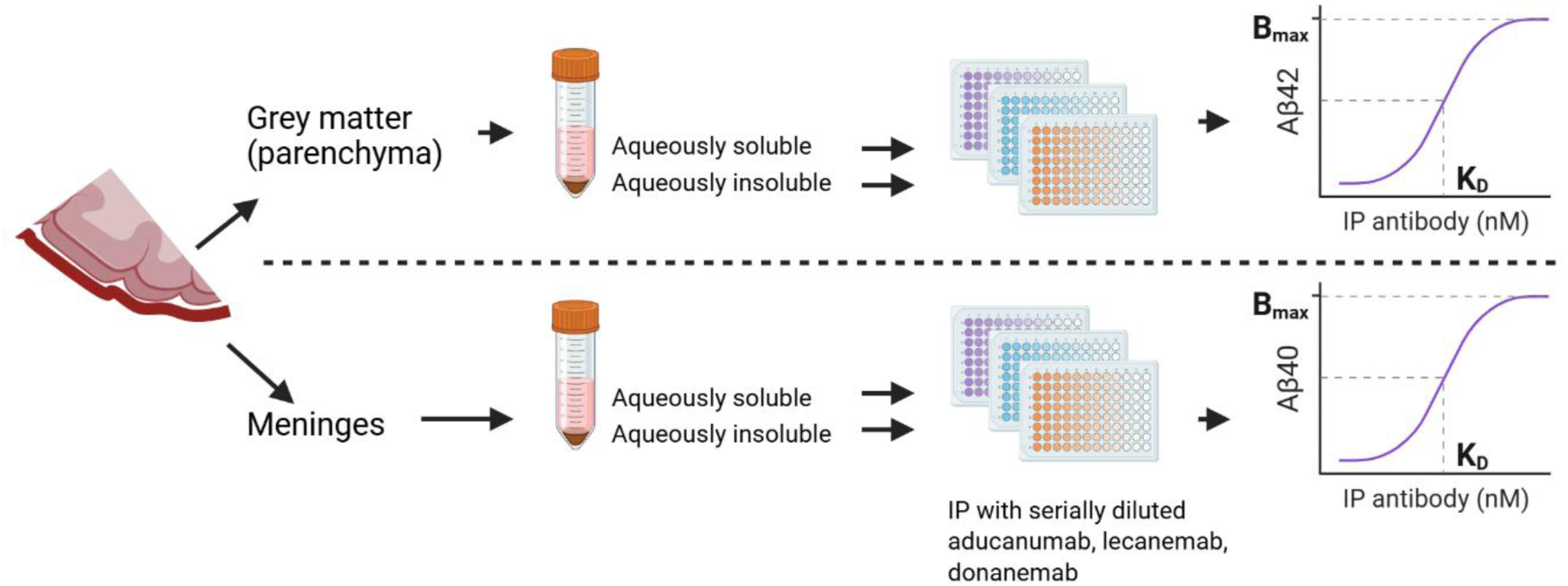
Experimental overview. Occipital lobe grey matter and overlying meninges from 18 cases with AD and CAA were processed to extract aqueously soluble and insoluble fractions. These were immunoprecipitated with serially diluted anti-amyloid antibodies followed by wash, elution, denaturation into monomers, and quantitation of total Aβ_42_ (for parenchyma) or Aβ_40_ (for meninges). Binding curves were fit to a one-site specific model and generated a K_D_, expressed in nM antibody and approximating equilibrium binding affinity, and a B_max_, expressed in ng/ml Aβ and reflecting the total amount of Aβ accessible to the antibody.

### Statistics

For our primary statistical analyses, we used the ratios of meningeal:parenchymal or insoluble:aqueous K_D_s and B_max_s as dependent variables. Because different antibody preparations can have different degrees of degradation or inactivity, normalizing by within-antibody ratios avoids this artifact. Expressing the K_D_ as a ratio can be interpreted as an antibody’s preference for one type of Aβ over another (*e.g.* soluble *vs* insoluble). We treated the unadjusted K_D_ and B_max_ without ratios as secondary outcomes. We refer to the meningeal:parenchymal ratios as “MP ratio” and the insoluble:soluble ratio as “IS ratio.”

All ratios were log-transformed for statistical analyses. We used a linear mixed model of the log(ratio) as the dependent variable, the random effect of patient (case), and the fixed effects of age, sex, antibody, and number of *APOE ε4* alleles. In all cases, the mixed models were fit by restricted maximum likelihood. T-tests used Satterthwaite’s method setting alpha to 0.05. Statistics were performed using R package lme4.

## Results

### Lecanemab does not exhibit greater preference for aqueously soluble (“protofibrillar”) Aβ compared to aducanumab

The definition of Aβ protofibrils extracted from AD brain has been those which remain in the supernatant after ultracentrifugation in aqueous buffer^5–9^. If lecanemab had greater preference for protofibrils than fibrils compared to other antibodies, then one would expect a higher ratio of its K_D_ for insoluble to soluble aggregates (IS K_D_ ratio) compared to other antibodies, in particular for parenchymal samples, where all the intended antibody targets lie. However, we detected no statistically significant difference between the lecanemab and aducanumab parenchymal IS K_D_ ratios (Fig 2A). The 95% confidence interval for the difference between lecanemab and aducanumab log(IS K_D_ ratios) was −0.220 to 0.174, meaning our results are 95% confident that lecanemab has between a 1.66-fold lesser and 1.49-fold greater preference for aqueously extracted Aβ (*i.e.* protofibrils) than aducanumab (Fig 2B). Thus, we conclude that lecanemab is unlikely to have greater preference for aqueously soluble Aβ aggregates than does aducanumab. However, our results did suggest that lecanemab and aducanumab have greater preference for aqueously extracted Aβ compared to donanemab (lecanemab 95% CI 0.029 to 0.422, 1.07-fold to 2.64-fold; aducanumab 95% CI 0.052 to 0.445, 1.12- to 2.79-fold) (Fig. 2B). Full model results presented in the supplementary tables.

**Figure 2.**
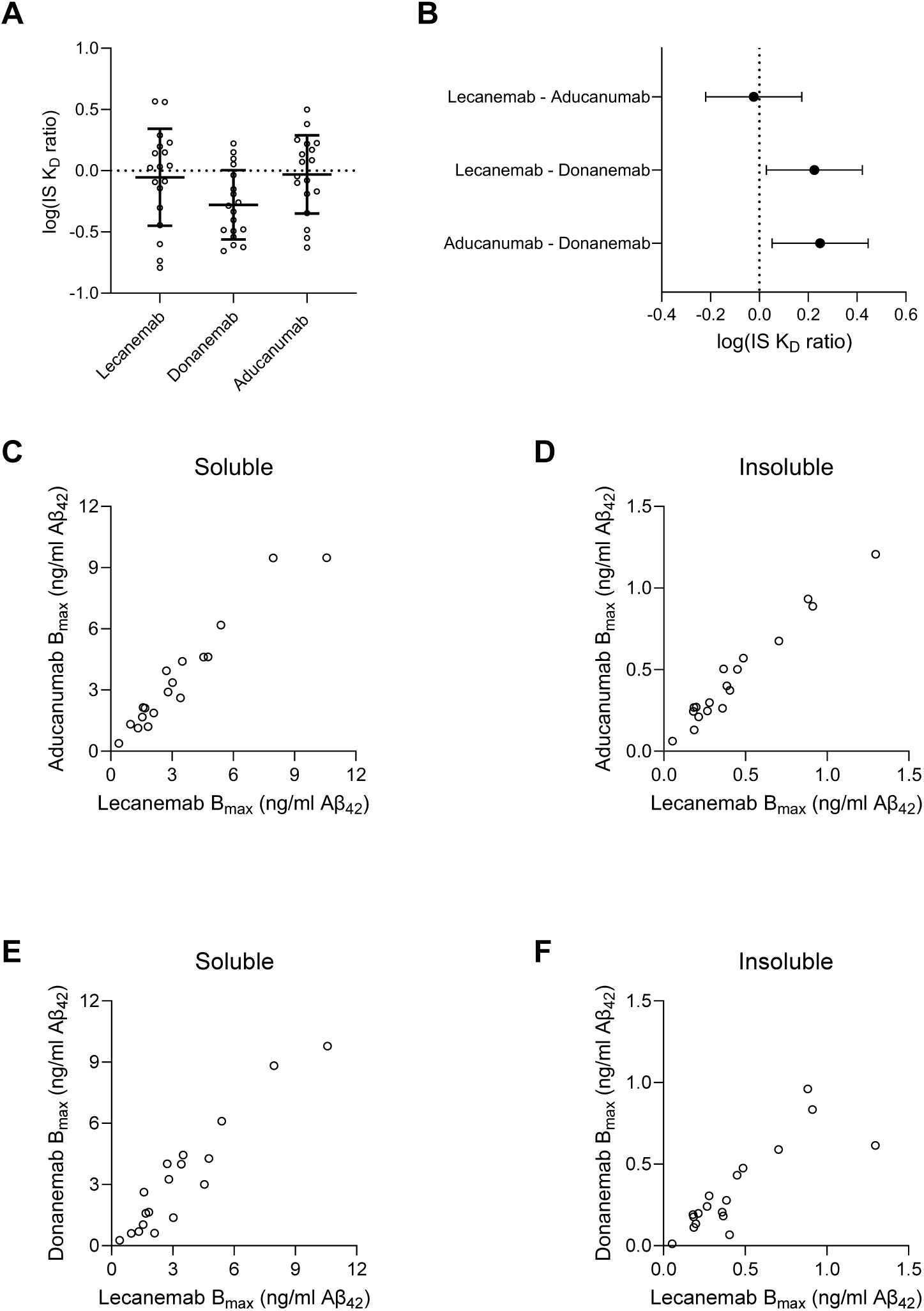
Binding profiles to insoluble *vs* soluble parenchymal Aβ_42_ aggregates. **(A)** The log(IS K_D_ ratio) reflects the binding preference of antibodies to insoluble *vs* soluble aggregates. A higher ratio implies greater preference for soluble aggregates. Donanemab exhibited a log ratio below 0, reflecting a slight preference for insoluble aggregates. Error bars = mean +/- SD. (**B)** Model estimates for the pairwise mean differences +/- 95% CI in log(IS K_D_ ratio). There was no statistically significant difference between lecanemab and aducanumab, but donanemab had a statistically different ratio compared to lecanemab and aducanumab. (**C-F)** Correlations between B_max_, the total Aβ accessible to the antibody, across soluble and insoluble extracts reveal near-perfect correlations.

### Lecanemab and aducanumab access the same pool of Aβ aggregates

The B_max_, the amount of Aβ immunoprecipitated at saturating antibody concentrations, is a measure of the total Aβ in a brain extract accessible to an antibody. If lecanemab bound a distinct pool of Aβ aggregates to which aducanumab or donanemab could not bind, then one might expect the B_max_ of lecanemab to exceed that of the others. Alternatively, if different AD patients’ brains contained different amounts of protofibrillar *vs.* fibrillar Aβ, then there would be an imperfect correlation between B_max_s of different antibodies across different cases. However, we found a perfect 1:1 correlation of lecanemab and aducanumab B_max_ in both the aqueously soluble and insoluble fractions (r = 0.97, P = 6.5E-11) (Fig 2C, D). The donanemab B_max_ also correlated with the B_max_ of lecanemab for soluble and insoluble fractions albeit with slightly weaker correlations (soluble r = 0.94, P = 4.5E-9; insoluble r = 0.84, p = 1.1E-5, Fig 2F). Overall, we conclude that all three antibodies access essentially identical populations of aggregates. Donanemab may bind only a subset of pyroglutamate-3 epitopes, but we reason that these are distributed evenly enough among the Aβ aggregates that all aggregates can be bound by donanemab at least one site.

### Antibody binding preference does not explain ARIA-E rates in clinical trials

In phase 3 trials, aducanumab caused ARIA-E in 35.2% of subjects receiving drug^10^, followed by 24.0% by donanemab^11^ and 12.6% by lecanemab^12^. If the differences in ARIA-E rates were due to differences in binding preferences for CAA vs plaque Aβ, then one would expect the ratio of K_D_ for meningeal Aβ_40_-rich to parenchymal Aβ_42_-rich aggregates (MP K_D_ ratio) to differ among the three antibodies and follow the order lecanemab > donanemab > aducanumab. We detected no statistically significant differences among the MP K_D_ ratios of the three antibodies in the soluble pool (Fig 3A). The 95% confidence interval for the difference in MP K_D_ ratio of lecanemab *vs.* aducanumab in the soluble pool was −0.115 to 0.291 (Fig 3B). Thus we conclude that in aqueous extracts, lecanemab has between a 1.30-fold lesser and a 1.95-fold greater preference for plaque *vs* CAA fibrils compared to aducanumab, unlikely to explain fully the ∼2.8-fold difference in ARIA-E rates in phase 3 clinical trials. Full model results are presented in the supplementary tables.

**Fig 3.**
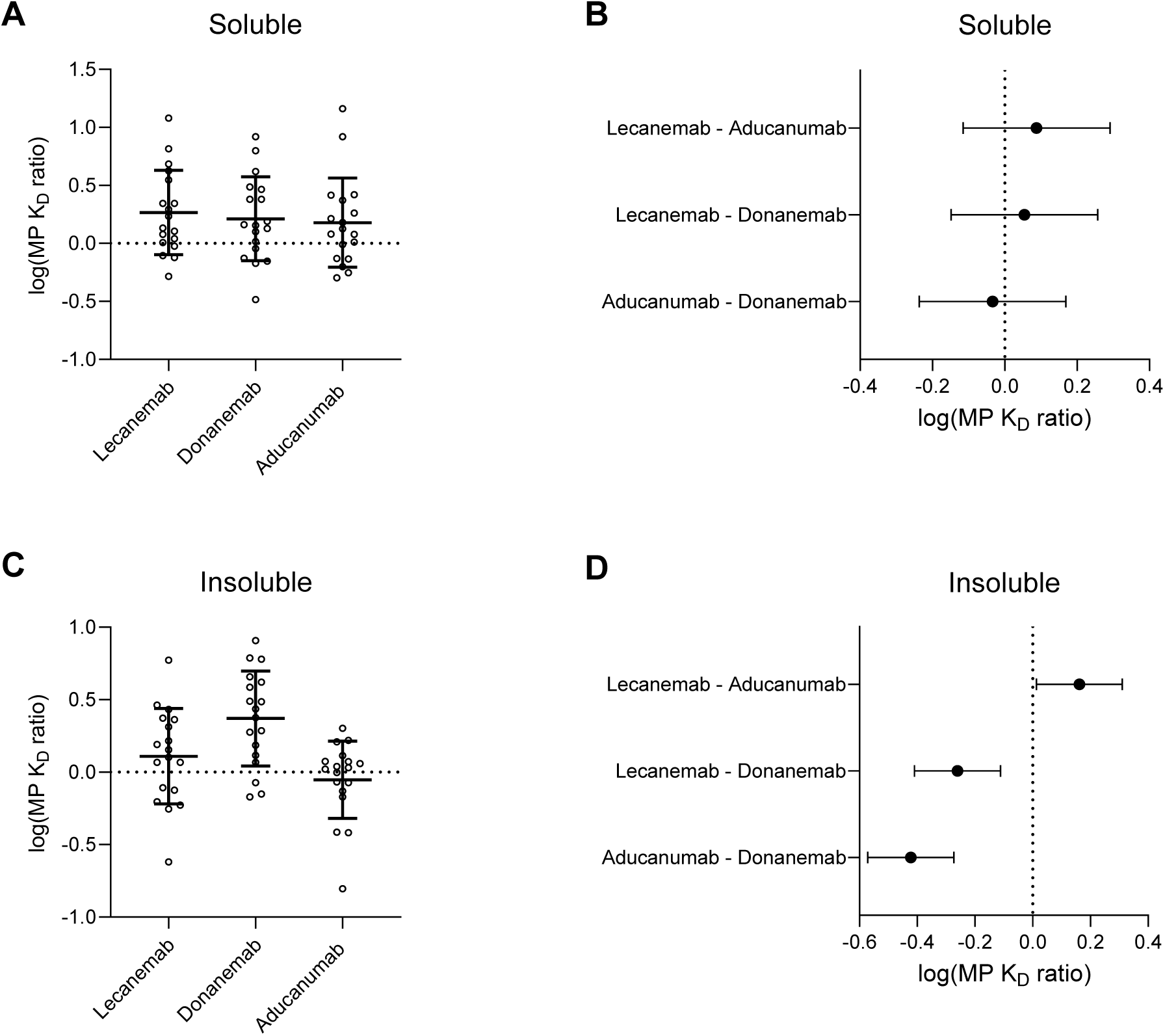
Plaque *vs* CAA antibody binding preferences. **(A, C)** The log(MP K_D_ ratio) reflects binding preferences of antibodies to meningeal Aβ_40_-rich aggregates (CAA-enriched) *vs* parenchymal Aβ_42_-rich aggregates (plaque-enriched). A higher ratio implies greater preference for plaque *vs* CAA aggregates. Error bars = mean +/- SD. **(B, D)** Model estimates for the pairwise mean differences +/- 95% CI in log(MP K_D_ ratio) reveals no significant differences in the soluble fraction but significant differences in the insoluble fraction. The difference in insoluble log(MP K_D_ ratio) between lecanemab and aducanumab reflects a 1.03- to 2.05-fold greater lecanemab preference for plaque compared to aducanumab, less than the ∼2.8-fold difference in phase 3 clinical trials.

In insoluble extracts, there were statistically significant differences in MP K_D_ ratio, including all three pairwise comparisons (Fig 3C). However, the rank order was donanemab > lecanemab > aducanumab, as opposed to the expected order lecanemab > donanemab > aducanumab based on phase 3 trial ARIA results. Examining only lecanemab and aducanumab, which do exhibit the hypothesized order based on trial results (lecanemab>aducanumab), the 95% confidence interval for the difference in MP K_D_ ratio was 0.013 to 0.311 (Fig 3D), meaning lecanemab exhibited between a 1.03- and 2.05-fold greater preference for plaque *vs* CAA fibrils compared to aducanumab. Although this difference is statistically significant, it may not fully explain the observed nearly 3-fold difference in ARIA rates, nor account for donanemab having still greater preference for plaque compared to lecanemab or aducanumab. Full model results are presented in the supplementary tables.

### Effect of *APOE* genotype on antibody binding

The *APOE ε4* allele increases the risk of Alzheimer disease through multiple mechanisms, including an increase in amyloid plaques, CAA, and in plaque-mediated tangle accumulation^13–16^. The *APOE ε4* allele also increases ARIA risk in a dose-dependent manner, and carriers may also benefit less from lecanemab and donanemab compared to non-carriers. We explored the effect of *APOE ε4* gene dosage in our antibody binding data to help explain these phenomena. We found the *APOE ε4* allele lowered the parenchymal IS B_max_ ratio in a dose-dependent manner for all antibodies; in other words, the *APOE ε4/ε4* homozygotes possessed a more soluble pool of Aβ accessible to each anti-amyloid antibody compared to heterozygotes, in turn more soluble than non-carriers (Fig 4A). In meninges, homozygotes possessed a lower IS B_max_ ratio than heterozygotes and non-carriers, but there was no difference between heterozygotes and non-carriers (Fig 4B). There was no statistically significant effect of *APOE ε4* gene dosage on the IS K_D_ ratio in either parenchyma or in meninges; in other words, *APOE ε4* did not affect the affinity with which antibodies bound to soluble *vs.* insoluble Aβ, only their total availability to bind. We also detected no effect of *APOE ε4* dosage on either the absolute K_D_ to meningeal extracts or those normalized to parenchymal extracts (the MP K_D_ ratio); in other words, *APOE ε4* did not change Aβ conformation in such a way to significantly alter antibody affinity. Full model results are presented in the supplementary tables.

**Fig 4.**
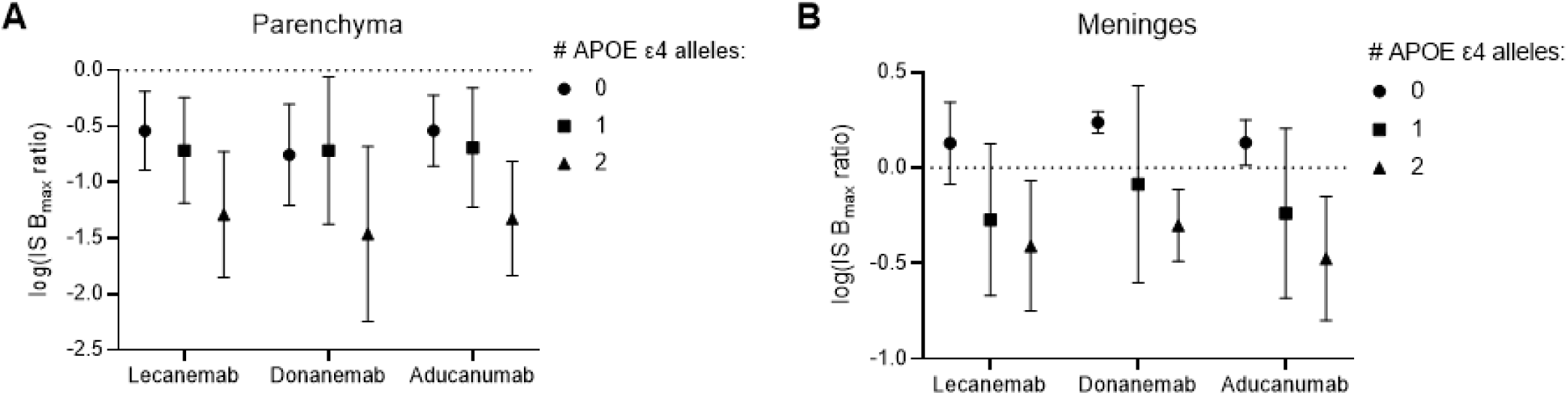
*APOE* genotype effects on Aβ accessible to antibody binding. The log(IS B_max_ ratio) reflects the solubility of the Aβ pool accessible to the antibody. A higher log(IS B_max_ ratio) reflects a less soluble pool of Aβ. We found that the *APOE ε4* dosage increased the solubility (decreased the log(IS B_max_ ratio)) of both parenchymal Aβ_42_ **(A)** and meningeal Aβ_40_ antibody targets. Error bars = mean +/- SD.

## Discussion

This study has important limitations. First, we only measured equilibrium binding constants (K_D_) as an approximation of affinity. We were unable to measure association (k_on_) and dissociation (k_off_) constants. Antibodies with identical equilibrium binding affinities can have different kinetics, and thus we cannot exclude that association and dissociation rates could differ among antibodies for parenchymal, vascular, soluble, or insoluble Aβ aggregates. We are developing techniques to measure these kinetics in AD brain extracts.

Second, the extraction method from the parenchyma could not separate plaque Aβ aggregates from microvascular CAA aggregates. Thus, some contamination in the parenchymal preparation with CAA aggregates was likely. However, we could enrich for plaque aggregates through the selective detection of Aβ_42_ over Aβ_40_ in the parenchymal fraction. Further, the meningeal preparations were unlikely to contain any plaque aggregates because amyloid plaques do not occur in the meninges. Thus, at best, the calculated MP ratios are of binding affinity to CAA *vs* plaque aggregates, and at worst, a ratio of binding affinity to CAA *vs* total (CAA + plaque) aggregates. This also assumes that CAA aggregates in meningeal blood vessels are biochemically equivalent to those in the parenchymal microvasculature, which they may not be. A last disadvantage of our approach is the exclusion of Aβ_40_-rich aggregates from the parenchyma; certain plaques are Aβ_40_-rich^17^, and antibody binding to these would not have contributed to our measures.

Third, we used antibodies produced recombinantly from patent sequences. While at least aducanumab and donanemab displayed similar binding in a subset of extracts (Fig S1), we cannot exclude subtle differences in binding compared to the brand-name product given to patients.

Fourth, we cannot exclude the extraction method itself causing structural alterations in the antibody epitopes. We have particular caution for interpreting donanemab results, because this is a non-conformational (linear) epitope. Homogenization may have sheared free pyroglutamate-3 Aβ monomers off of aggregates, leading to an excess of monomeric pyroglutamate-3 Aβ as opposed to that which naturally occurs in the aggregates. This artificially sheared pyroglutamate-3 Aβ could result in falsely low K_D_ and B_max_ values. Because aducanumab and lecanemab recognize conformational epitopes, shearing of monomers should not have caused this artifact, though we cannot exclude alterations in their epitope abundance during extraction.

Within these limitations, we conclude that 1) lecanemab does not bind a distinct population of Aβ aggregates inaccessible to aducanumab or donanemab, 2) that lecanemab exhibits equivalent binding affinity to “soluble protofibrils” (aqueously extracted Aβ aggregates) than aducanumab or donanemab, and 3) that the observation from phase 3 trials that lecanemab caused less ARIA-E than donanemab, in turn less than aducanumab, can be explained by differences in binding preference to CAA. In previous work, we found that anti-amyloid antibodies including lecanemab could bind to aqueously extracted short Aβ fibrils from human brain^18^, suggesting that much of what has previously been termed “protofibrils” and presumed to possess a different structure (and thus different antibody binding characteristics) compared to fibrils, are in fact structurally the same amyloid fibrils as those found in amyloid plaques. Our data from this study are consistent with this model: a putatively protofibril-preferring antibody (lecanemab) exhibited the same binding preference to the aqueous (“protofibrillar”) extraction as did the putatively non-protofibril-preferring antibody (aducanumab). Previous comparisons of lecanemab to other antibodies used *in vitro* synthetic preparations of Aβ^1^, the structures of which are unknown and may not exist in Alzheimer disease. One previous measurement comparing lecanemab to other antibodies’ binding to human meningeal extracts did not include a ratio to parenchymal extracts, tested only N=4 brains, and excluded one of these as an outlier^19^.

Our results do not exclude off-target CAA-binding as a cause of ARIA-E. Rather, we conclude only that *differences* between antibodies’ ARIA-E rates and efficacy cannot be explained by *differences* in the equilibrium binding constants we measured. A small dose titration adjustment in the Trailblazer-Alz 6 trial of donanemab reduced ARIA rates^20^, implying that at least some differences in ARIA-E rates to other antibodies could be explained by dosing or monitoring factors rather than differences in antibody affinity. The phase 3 trials for aducanumab, donanemab, and lecanemab differed in titration regimens and in the number of monitoring MRIs obtained; the latter could lead to sampling bias. While all three antibodies are IgG1 subclass, differences in Fc receptor engagement, half-life, off-rates or other antibody-dependent but binding equilibrium affinity-independent factors could additionally affect ARIA-E and efficacy.

We found that the *APOE ε4* allele was associated with an overall more soluble pool of Aβ that the antibodies could bind to. Future studies may relate how this alteration may affect therapeutic parameters such as clearance from the brain and neutralization of toxicity, and whether a more soluble pool of CAA or plaque Aβ results in an elevated risk of ARIA-E.

Future efforts to improve anti-amyloid therapy for Alzheimer disease will require a deeper mechanistic understanding of how antibody-antigen interactions shape plaque clearance, clinical efficacy, and ARIA.

## Acknowledgements

We thank Reisa Sperling for her valuable insights. This work was funded by National Institutes of Health awards K08NS128329 (Stern) and R01NS136122 (Lemere, Stern), and the Davis Alzheimer Prevention Program (Selkoe). We thank the NeuroTechnology Studio at Brigham and Women’s Hospital for providing funding for brain banking efforts. DJS is a director of Prothena Biosciences and ad hoc consultant to Roche and Eisai. CAL is a consultant or scientific advisory board member to Acumen Pharmaceuticals, Advantage Therapeutics, Apellis Pharmaceuticals, Cyclotherapeutics, Eli Lilly, Merck, MindImmune, Novo Nordisk, Sanofi, Switch Therapeutics, and Takeda Pharmaceuticals. The other authors declare no conflicts of interest.

## Supplementary Figures

**Fig S1.**
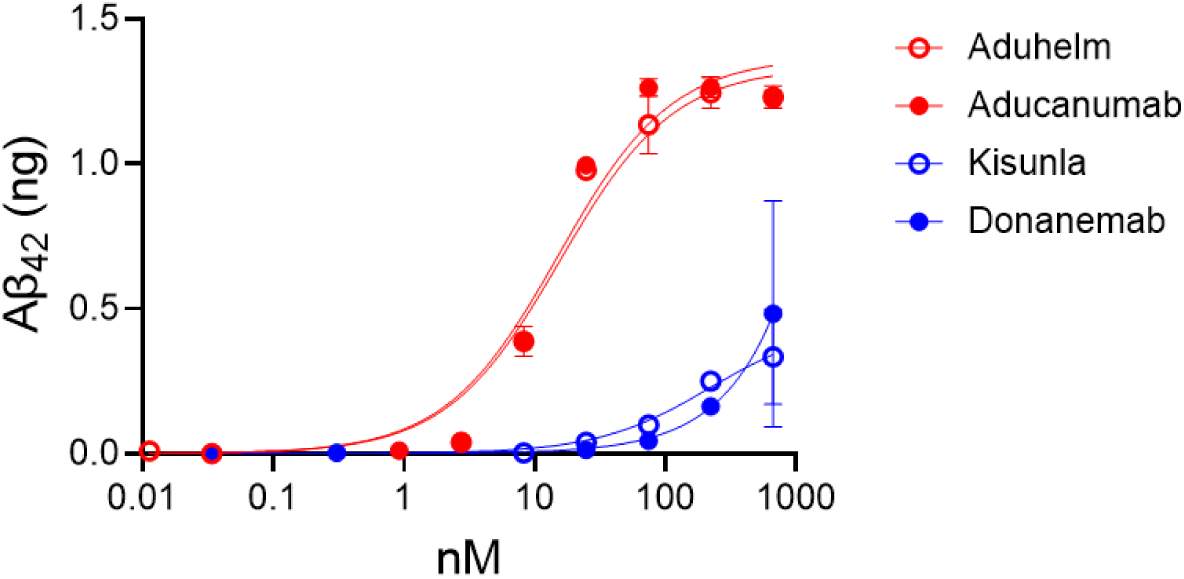
Binding profile of recombinant aducanumab and donanemab equivalents used for this study were identical to those of their brand-name products.

**Fig S2.**
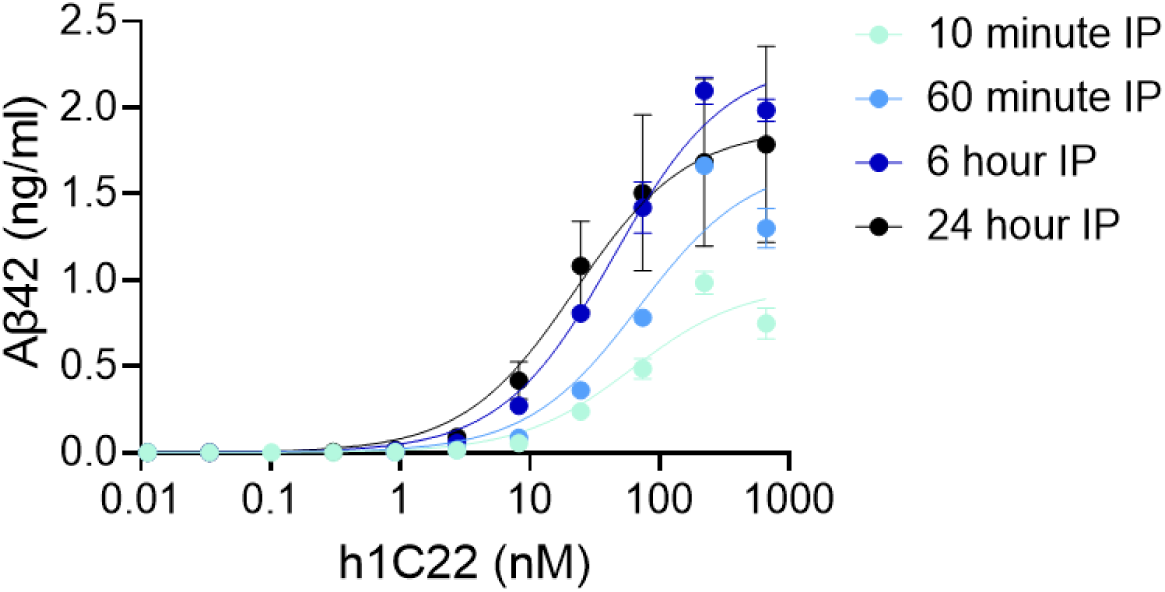
Past 6 hours, the binding curve for test antibody h1C22 equilibrated. Thus, we used a 24-hour immunoprecipitation reaction to measure equilibrium binding of anti-amyloid antibodies.

**Fig S3.**
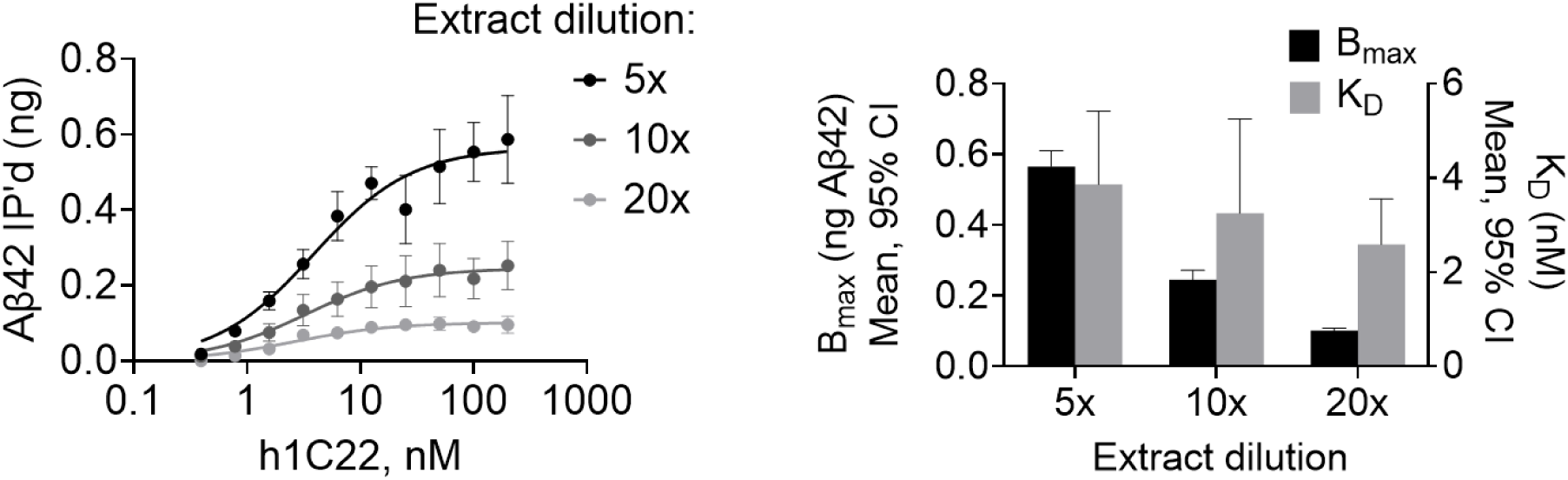
As a human AD brain extracted is diluted 5-, 10-, or 20-fold, the B_max_ diminishes proportionately to the dilution factor but the K_D_ does not, implying a measurement of antibody affinity rather than antibody titration.

**Fig S4.**
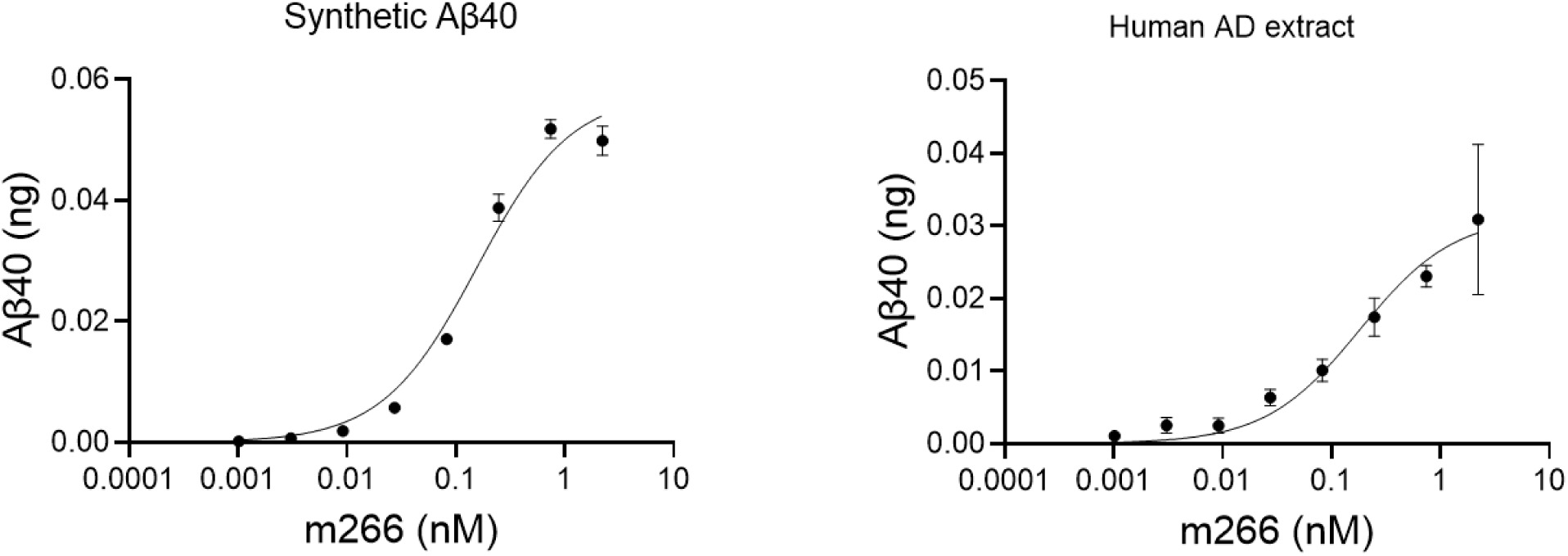
The binding of test antibody m266, which binds the Aβ mid-region and selectively binds monomers, exhibits the same binding profile with synthetic Aβ_40_ monomers as those found naturally in a human AD extract. This antibody was chosen to test the validity of the method because the m266 epitope, being linear and present on monomers, is expected to have the same structure in synthetic as human brain-derived extracts.

## Supplementary Methods

### Extraction of Aβ aggregates from parenchyma and meninges

We optimized a protocol based on multiple literature sources^21–23^ for the extraction of aqueous and insoluble Aβ aggregates from both grey matter and meninges. First, an occipital coronal slice of frozen brain tissue was thawed on ice. The leptomeninges were peeled off with forceps and then chopped on a McIlwain tissue chopper set to 0.05 mM. The grey matter was dissected from the white matter with a scalpel. Both the grey matter and meninges were weighed and resuspended five volumes TBS extraction buffer containing 25 mM Tris, 150 mM NaCl, pH 7.4 supplemented with 5 μg/ml leupeptin, 5 μg/ml aprotinin, 2 μg/ml pepstatin, 120 μg/ml 4-benzenesulfonyl fluoride hydrochloride, and 5 mM NaF. Both were homogenized using a Teflon Dounce homogenizer attached to an IKA stirrer at 800 rpm for 25 strokes. The parenchymal homogenate, being of greater volume, was centrifuged in a Type 70.1Ti rotor (Beckman) in thickwall polycarbonate tubes (Beckman cat. 355630) at 25,000 rpm (rcf_av_ = 42,800 *g*) for 20 min at 4°C. The meningeal homogenate was centrifuged in 1.5-ml microcentrifuge tubes in a tabletop centrifuge at 20,000 *g* for 2 hours. We found that these two centrifugation protocols, adapted to different volumes, resulted in similar amounts of Aβ in the supernatant for the same sample. The top 90% supernatant was retained as the aqueously extracted Aβ fraction, and the bottom 10% discarded. The pellets were frozen at −80°C for later processing.

To extract insoluble Aβ from parenchyma, the parenchymal pellet was thawed, re-weighed, and resuspended in five volumes TBS extraction buffer supplemented with 2% N-lauryl sarcosine (sarkosyl). The suspension was incubated at 37°C for 1 hour, then re-homogenized as above. The homogenate was centrifuged in a TLA-55 rotor at 55,000 rpm (rcf_av_ = 136,000 x *g*) for 1 hour at 4°C. The pellet was washed twice in 1 ml TBS without sarkosyl by resuspending and centrifuging again in a TLA-55 rotor at 55,000 rpm (rcf_av_ = 136,000 x *g*) for 1 hour at 4°C. The final pellet was weighed once more and resuspended in five volumes TBS, then sonicated for 25 seconds at 35% power prior to aliquoting. After thawing and dilution for IP-ELISA, the extract was sonicated again for 5 minutes at 35% power.

To generate insoluble Aβ from meninges, we followed a protocol used in the literature to derive meningeal fibrils^22,24^. The meningeal pellet from the aqueous extraction was thawed and sliced again with a scalpel. The sliced material was weighed again and washed three times in five volumes Tris-calcium buffer containing 20 mM Tris, pH 8.0 with 138 mM NaCl, 2 mM CaCl_2_, and 0.1% NaN_3_ followed by centrifuging at 20,000 *g* for 5 minutes in a tabletop centrifuge. The washed pellet was then resuspended once more in Tris-calcium buffer supplemented with 5 mg/ml collagenase from *Clostridium histolyticum* (Sigma) and nutated overnight at 37°C. Remaining steps were carried out on ice or at 4°C. The suspension was centrifuged at 20,000 *g* for 10 minutes in a tabletop centrifuge. The pellet was then washed twice by resuspending in two volumes wash buffer containing 50 mM Tris, 10 mM EDTA, pH 8.0 and centrifuging in a TLA-55 rotor at 55,000 rpm (rcf_av_ = 136,000 x *g*). The pellet was resuspended in one volume deionized water and homogenized with a hand-held homogenizer (Fisher), then centrifuged at 12,000 *g* for 5 minutes. The supernatant was saved, and the pellet was again resuspended in water and centrifuged again at 12,000 *g* for 5 minutes. This water extraction was performed five times, and all five fractions were pooled and then aliquoted for IP-ELISA.

### Antibody sources

Recombinant human aducanumab, lecanemab, and donanemab were produced recombinantly by Evitria, Inc. in Chinese Hamster Ovary (CHO) cells according to the amino acid sequence found in the patent literature. Antibody purity was verified by HPLC. Mouse antibodies 266, 2G3 and 21F12 were gifts from Elan Pharmaceuticals. Aducanumab-avwa (Aduhelm) and donanemab-azbt (Kisunla) for verification experiments were purchased from McKesson.

### Measurement of antibody binding to Aβ aggregates from human brain

Brain extracts were diluted in TBS-T (25 mM Tris pH 7.4, 500 mM NaCl, 0.05% Tween-20) in the presence of 20 µl washed protein G beads (Bio-Rad) and a three-fold dilution series of the IP antibody starting at 100 µg/ml (666.67 nM) in 96-well round-bottom protein low-binding plates (Greiner), in a final volume of 150 µl. To approximate the same concentrations of antigen, aqueous extracts were diluted 10-fold, insoluble parenchymal extracts were diluted 2,000-fold, and insoluble meningeal extracts were diluted 200-fold. The IP reactions were carried out at 4°C for 18 hours on a plate shaker at 800 rpm. The beads were washed three times in 200 µl TBS-T followed by elution in 50 µl 5 M GuHCl shaking overnight at 4°C. The eluate was then quantified by MSD Aβ ELISAs (see method below). The Aβ42 assay was used to quantify the parenchymal IP, and the Aβ40 assay to quantify the meningeal IP. The IP antibody concentration in nM was normalized to the background (0 nM antibody) by subtraction and plotted against the amount of Aβ IP’d. The data were fitted to a one-site specific binding model (GraphPad Prism) of the form Aβ = B_max_*X/(K_D_ + X), where X represents the IP antibody concentration. Fitting was accomplished by least squares regression without weighting or constraints. We regard the K_D_ as an approximate measure of binding affinity, and the B_max_ as a measure of the total Aβ available for binding by the test antibody. The K_D_ is not a precise measure of affinity, however, because the molar quantity of Aβ aggregates is unknown – only the concentration of total monomers composing those aggregates is quantitated. However, by comparing all antibodies across all brains, we can make relative comparisons between antibodies, even if the K_D_ is not an exact measure of affinity.

We used the excellent paper by Jarmoskaite *et al.*^25^ as a guide to ensure we measured binding properties of the antibodies of interest. We validated the method using a humanized version of test antibody 1C22, for which *in vitro* binding characteristics had previously been evaluated^26^. Past six hours of incubation time, the binding curves had saturated (Fig S2), indicating the reaction was at equilibrium – we chose an 18-hour binding time going forward for convenience. We serially diluted the brain extracts and evaluated the binding curves (Fig S3), finding that the B_max_ was proportionately lowered as expected, but the K_D_, was not, indicating that the K_D_ is measuring a binding property of the antibody-antigen interaction, rather than a simple titration of the antibody out of solution as would be the case if the antigen concentration were too high^25^.

We tested whether the IP-ELISA method estimated the binding affinities similarly in complex brain extracts as in pure solution. Using IP antibody m266, which has strong preference for monomers, we evaluated its K_D_ in pure solution for Aβ40 (Fig S4) at approximately 23 nM, which agreed with the value for a meningeal AD extract at 22 nM.

We were able to purchase brand-name Aduhelm (aducanumab-avwa) and Kisunla (donanemab-azbt), and compare these binding properties to the recombinant aducanumab and donanemab used in this study (Fig S1), showing no significant difference.

### MSD ELISAs

Aβ monomer-preferring ELISAs were home-brew immunoassays as described^18^. Samples containing GuHCl were diluted such that the GuHCl concentration was ≤0.25 M to avoid interference with the antibody reaction. All steps were at room temperature. Plates were coated with capture antibody 266 at 3 µg/ml in PBS overnight and blocked in 5% MSD Blocker A in TBS with 0.05% Tween-20 (TBS-T) for 1.5 h. Samples were then applied for 1.5 h after being diluted in 1% Blocker A in TBS-T. Plates were washed 3 times in TBS-T, then biotinylated detector antibody and MSD Strepavidin-Sulfotag (1:5,000) were applied for 1.5 h. Biotinylated antibodies 21F12 at 0.4 µg/ml and 2G3 at 0.2 µg/ml were used as detectors for the Aβ42 and Aβ40 assays, respectively. After three more washes in TBS-T, the plates were detected with 2x MSD read buffer T. The lower limit of quantification (LLoQ) was defined as the lowest standard with a luminescence value at least twice the blank average, and the lower limit of detection (LLoD) was defined as the lowest standard with values greater than the blank average plus twice the blank standard deviation.

## Supplementary Tables

**Table S1.** Fixed effects of linear mixed model for parenchymal log(IS K_D_ ratio).

|  | Estimate | Std. Error | df | t value | Pr(>\|t\|) |
| --- | --- | --- | --- | --- | --- |
| (Intercept) | -0.17825 | 0.879401 | 14.07809 | -0.20269 | 0.842274 |
| Aducanumab | 0.023238 | 0.080296 | 34 | 0.289406 | 0.774028 |
| Donanemab | -0.22528 | 0.080296 | 34 | -2.80563 | 0.008248 |
| APOE (# ε4) | 0.193307 | 0.090766 | 14 | 2.129725 | 0.05142 |
| Age | -0.00352 | 0.012026 | 14 | -0.29284 | 0.773939 |
| Female sex | 0.321972 | 0.144682 | 14 | 2.225369 | 0.043003 |

**Table S2.** Fixed effects of linear mixed model for soluble log(MP K_D_ ratio).

|  | Estimate | Std. Error | df | t value | Pr(>\|t\|) |
| --- | --- | --- | --- | --- | --- |
| (Intercept) | -0.49904 | 1.083065 | 14.05464 | -0.46077 | 0.652014 |
| Aducanumab | -0.0879 | 0.082765 | 34 | -1.06207 | 0.295689 |
| Donanemab | -0.05415 | 0.082765 | 34 | -0.65432 | 0.517308 |
| APOE (# ε4) | 0.222131 | 0.111834 | 14 | 1.986261 | 0.066942 |
| Age | 0.006821 | 0.014817 | 14 | 0.460373 | 0.652318 |
| Female sex | -0.00129 | 0.178264 | 14 | -0.00725 | 0.994314 |

**Table S3.** Fixed effects of linear mixed model for insoluble log(MP K_D_ ratio).

|  | Estimate | Std. Error | df | t value | Pr(>\|t\|) |
| --- | --- | --- | --- | --- | --- |
| (Intercept) | 0.145335 | 1.043693 | 14.03173 | 0.139251 | 0.891231 |
| Aducanumab | -0.16181 | 0.060808 | 34 | -2.66098 | 0.011806 |
| Donanemab | 0.26036 | 0.060808 | 34 | 4.281659 | 0.000143 |
| APOE (# ε4) | -0.04043 | 0.107812 | 14 | -0.37499 | 0.713285 |
| Age | 0.001281 | 0.014284 | 14 | 0.089665 | 0.929824 |
| Female sex | -0.16735 | 0.171854 | 14 | -0.97377 | 0.346697 |

**Table S4.**
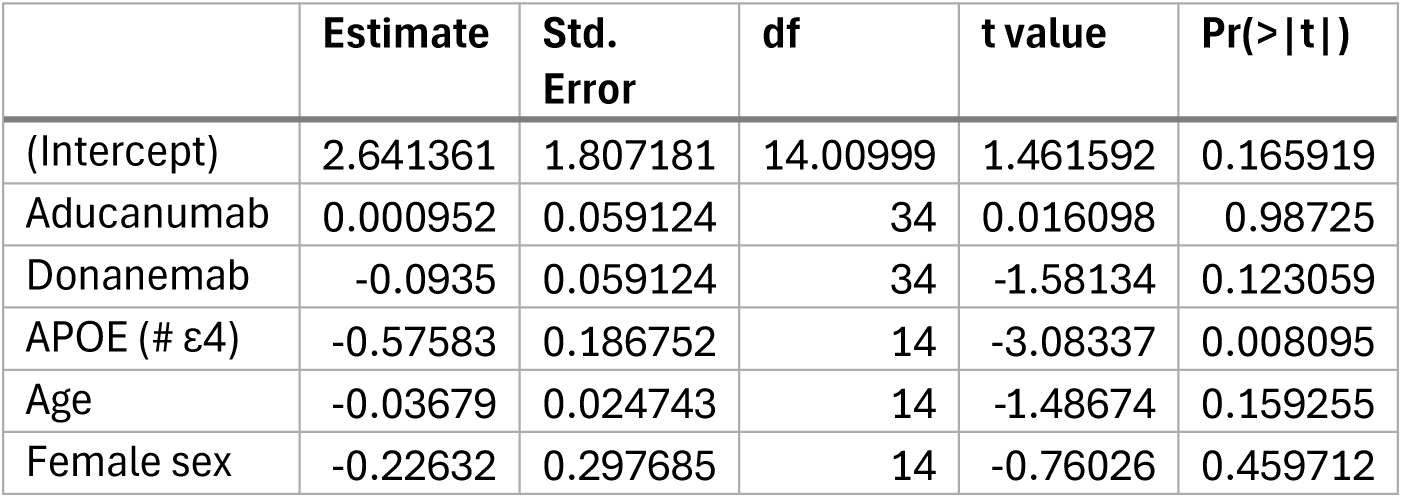
Fixed effects of linear mixed model for parenchymal log(IS B_max_ ratio).

